# Sustained efficacy of CRISPR-Cas13b gene therapy for FSHD is challenged by immune response to Cas13b

**DOI:** 10.1101/2024.12.18.629250

**Authors:** Afrooz Rashnonejad, Manal Farea, Gholamhossein Amini-Chermahini, Gerald Coulis, Noah Taylor, Allison Fowler, Armando Villalta, Oliver D. King, Scott Q. Harper

## Abstract

Facioscapulohumeral muscular dystrophy (FSHD) is a potentially devastating muscle disease caused by de-repression of the toxic *DUX4* gene in skeletal muscle. FSHD patients may benefit from *DUX4* inhibition therapies, and although several experimental strategies to reduce *DUX4* levels in skeletal muscle are being developed, no approved disease modifying therapies currently exist. We developed a CRISPR-Cas13b system that cleaves *DUX4* mRNA and reduces DUX4 protein level, protects cells from DUX4-mediated death, and reduces FSHD-associated biomarkers *in vitro*. *In vivo* delivery of the CRISPR-Cas13b system with adeno-associated viral vectors reduced acute damage caused by high *DUX4* levels in a mouse model of severe FSHD. However, protection was not sustained over time, with decreases in Cas13b and guide RNA levels between 8 weeks and 6 months after injection. In addition, wild-type mice injected with AAV6.Cas13b showed muscle inflammation with infiltrates containing Cas13b-responsive CD8+ cytotoxic T cells. Our RNA-seq data confirmed that several immune response pathways were significantly increased in human FSHD myoblasts transfected with Cas13b. Overall, our findings suggest that CRISPR-Cas13b is highly effective for *DUX4* silencing but successful implementation of CRISPR/Cas13-based gene therapies may require strategies to mitigate immune responses.

## Introduction

Facioscapulohumeral muscular dystrophy (FSHD) is among the most prevalent forms of muscular dystrophy, possibly affecting up to 1 in 8,000 individuals (*1, 2*). FSHD is historically described as a progressive myopathy associated with wasting and weakness in muscles of the face, shoulder girdle, and arms (*3*). Although this classical presentation occurs in many individuals, FSHD can be highly variable, even within families (*4-6*). Severely affected patients may develop widespread muscle weakness leading to ambulation problems, wheelchair dependence, requirement for caregiver assistance, and respiratory muscle involvement that can contribute to death, while other patients with similar FSHD-permissive genetics may be mildly affected or unaffected (*7*). Many patients report debilitating pain (*8, 9*). Age-at-onset and rate of progression are also variable, and asymmetry is common within affected individuals. Thus, non-uniform presentation is a hallmark of the disease (*4, 5, 10, 11*).

The search for an FSHD gene began with the emergence of positional cloning, but solving the complex and atypical pathogenic mechanisms underlying FSHD required decades of research. Several studies now support that FSHD is caused by aberrant expression of the *double homeodomain 4* (*DUX4*) gene in skeletal muscle (*12-24*). In short, *DUX4* is normally off in skeletal muscle, but epigenetically de-repressed in muscles of FSHD patients, leading to cell death and dysfunction (*12, 14, 17-19, 22*). The identification of *DUX4* as a primary FSHD gene enabled development of *DUX4*-expressing cell (*13, 25-30*) and animal models (*15, 19, 21, 31-42*), and provided a rational therapeutic target. We hypothesize that the most direct approach to FSHD therapy will involve *DUX4* inhibition, and numerous strategies are being developed to accomplish this goal at the DNA (*43-46*), RNA (*23, 47-57*) and protein levels (*58-60*).

In this study, we sought to test a new gene therapy strategy to inhibit *DUX4* mRNA using CRISPR-Cas13 technology. Like the more commonly used Cas9 protein, Cas13 is a bacteria-derived, RNA-guided CRISPR-associated nuclease. However, while Cas9 cuts DNA, Cas13 cleaves RNA (*61-64*). Since Cas13 does not cut DNA, we reasoned that its use could avoid possible safety concerns related to genome editing following long-term muscle expression of Cas9 from a viral vector. In addition, our rationale for pursuing Cas13-mediated FSHD gene therapy, versus one employing Cas9, was rooted in the genetics of FSHD. In particular, the human *DUX4* gene is embedded in 3,300 base pair repetitive elements called D4Z4 repeats (*65, 66*). Potentially hundreds of D4Z4/DUX4 copies exist in the human genome, on multiple chromosomes, but only one copy, located nearest the human chromosome 4q35 telomere, expresses a stabilized, polyadenylated *DUX4* mRNA in FSHD muscle (*14, 66*). Thus, a Cas13-associated guide RNA targeting the FSHD-permissive *DUX4* open reading frame (ORF) would direct silencing of *DUX4* mRNA arising from a single locus in FSHD muscle, while an analogous Cas9-associated guide RNA would direct Cas9 to cleave the human genome in potentially hundreds of places unless targeted to the poly(A) signal, which, although possible, has limited options for gRNA design (*45, 47, 67, 68*). We therefore reasoned that Cas13-mediated RNA editing could be better indicated for FSHD than an analogous Cas9-based DNA editing system.

At the inception of this study, there were only two Cas13 orthologs in published literature, Cas13a from *Leptotrichia shahii* (*61*) and Cas13b from *Prevotella sp. P5-125* (*62, 63*). We began with Cas13b because it was suggested to be more efficient and have less collateral activity (*62*). This system efficiently reduced *DUX4* expression in cultured human cells and mouse muscle, but *DUX4* silencing was not sustained *in vivo*. Here we report the first evidence that an immune response to Cas13 was likely associated with loss of silencing efficacy over time in an animal model. Our findings suggest successful implementation of CRISPR/Cas13-based gene therapies may require strategies to mitigate immune responses.

## RESULTS

### Design and In Vitro Efficacy Screening of Cas13b gRNAs Targeting DUX4 mRNA

In this study, we developed and evaluated a *DUX4* mRNA-targeting Cas13b system as a potential treatment for FSHD. First, we designed fifteen guide RNAs (gRNAs) targeting the *DUX4* coding and 3’ untranslated (UTR) regions (**Fig 1A-B**). In contrast to CRIPSR-Cas9 systems, RNA-cleaving Cas13 enzymes do not use guide RNAs with nucleotide protospacer adjacent (PAM) motifs (*14, 62, 66*). The lack of a requirement for PAM sequences suggested that Cas13 gRNAs could be designed against any stretch of sequence, but we were uncertain about how to best construct candidate gRNAs because no clear design rules were in place when we began this project. However, some data suggested Cas13 gRNAs more efficiently target unstructured, single stranded RNA regions (*61, 64*) and we therefore used the RNAfold program (http://rna.tbi.univie.ac.at/cgi-bin/RNAWebSuite/RNAfold.cgi) to predict *DUX4* mRNA secondary structure, and subsequently designed 15 gRNAs targeting putatively unstructured regions (*69-71*) (**Fig 1A-B**). We designed all gRNAs as DNA expression constructs containing 4 main sequence features: (1) a human U6 promoter to drive transcription; (2) a 36 nucleotide (nt) Cas13b-binding direct repeat at the 5′-end of the transcript; (3) 29-34 nucleotides of antisense homology to human *DUX4* mRNA; followed by (4) an RNA polymerase III termination signal (6 thymine nucleotides) (**Fig 1C, Table S1**) (*62, 63*).

**Fig 1.**
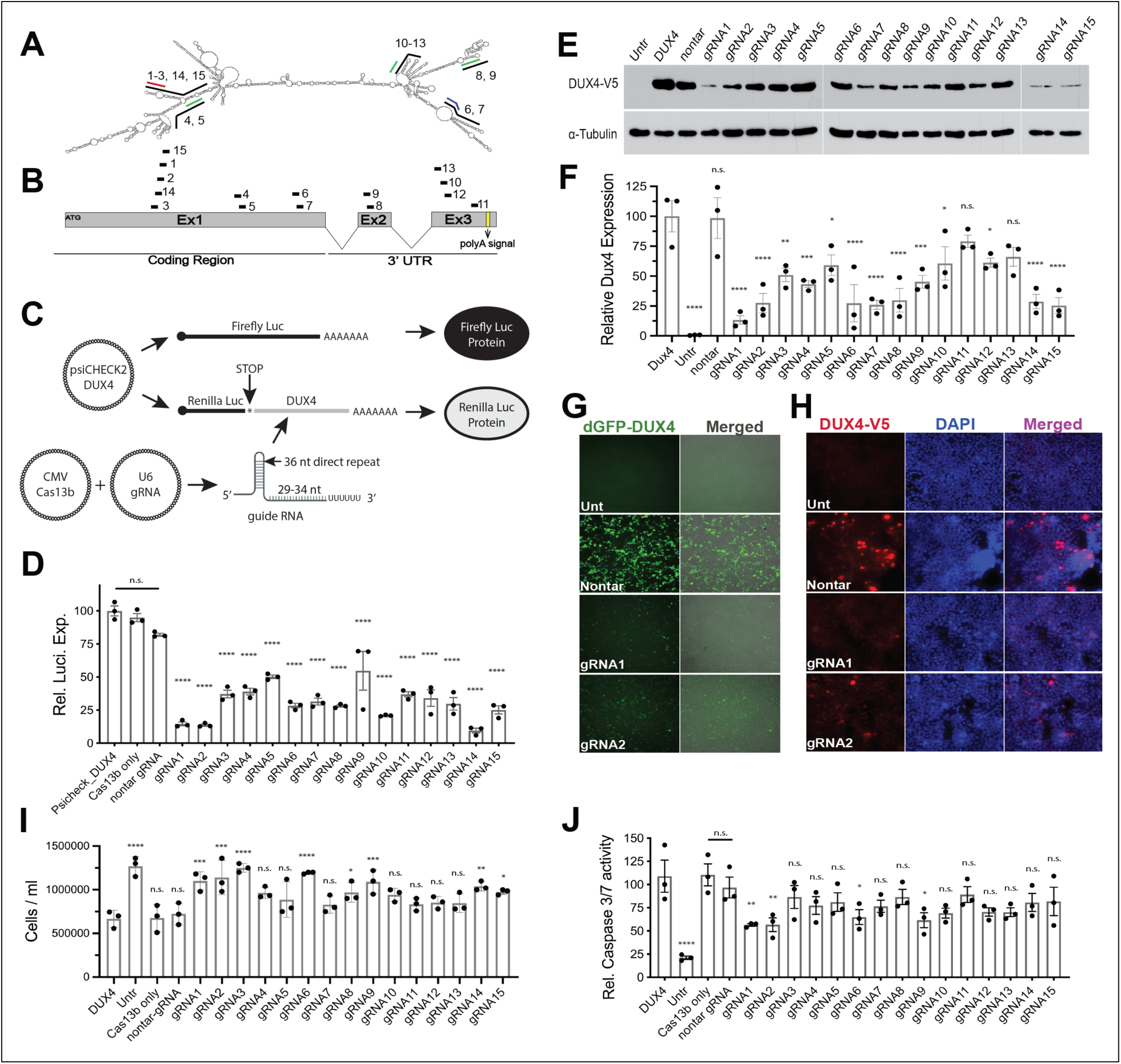
Cas13b gRNAs efficiently target DUX4 and protect cells from death. (A) *DUX4* mRNA secondary structure and gRNAs target sites. (B) Illustration of 15 gRNAs targeting different parts of the *DUX4* mRNA. (C) Schematic drawing of Luciferase assay using psiCHECK2-DUX4 Dual Luciferase system. DUX4-FL sequences are cloned as the 3’UTR of the *Renilla* luciferase gene, while Firefly luciferase serves as a transfection control in same plasmid. HEK293 cells were co-transfection with plasmids expressing Cas13b, each gRNA, and psiCHECK2-DUX4. Successful targeting of DUX4 sequences by Cas13b-gRNA complexes leads to reduction of *Renilla* luciferase expression. (D) All gRNAs significantly reduced *Renilla* luciferase-*DUX4* with gRNA1, gRNA2, gRNA6, gRNA7, gRNA8, gRNA10, gRNA14 and gRNA15 yielding the most efficient silencing. (E-F) Western blot results showing reduced DUX4 protein level in Cas13b-gRNA-treated cells compared to nontar-gRNA with gRNA1, gRNA2, gRNA6, gRNA7, gRNA8, gRNA9, gRNA14, and gRNA15 showing the most consistent silencing. In contrast, the non-targeting control did not impact DUX4 protien levels. α-Tubulin was used as endogenous loading control. (G) Using a destabilized GFP-DUX4 fusion protein (dGFP-DUX4), both lead gRNA 1 and gRNA2 efficiently reduced GFP signal compared to nontar-gRNA treated cells. 10X objective. Unt: untransfected cells. Merged: GFP images combined with black-and-white light microscopy images to show cell density. (H) Immunofluorescence showing reduced DUX4.V5 protein expression in HEK293 cells transfected with plasmids expressing V5-epitope tagged *DUX4*, Cas13b, and gRNAs 1 or 2, 24 hrs post-transfection. 20X objective. (I-J) DUX4 activates HEK293 cell death by 48 hrs post-transfection, which can be measured by cell viability or caspase-3/7 assays. (I) Viability improved significantly in cells treated with gRNA1, 2, 3, 6, 8, 9, and 14 (J), while only gRNA1 (56.4 ± 1.35%), gRNA2 (56.67 ± 7.41%), gRNA6 (64.86 ± 7.99%), and gRNA9 (61.59 ± 8.07%) significantly reduced caspase-3/7 activity. N=3 independent experiments for all studies. For luciferase, cell viability, and caspase 3/7 assay, each independent experiment was performed in triplicate. *P < 0.05, **P < 0.01, ***P < 0.001, and ****P < 0.0001, ANOVA.

We next performed primary, secondary and tertiary screens in HEK293 cells to empirically identify the most efficient *DUX4*-silencing gRNAs *in vitro*. We began with HEK293 cells because they rapidly divide, are easily transfected, and undergo DUX4-induced cell death. For a primary efficacy screen, we developed a dual luciferase assay using a plasmid containing full-length *DUX4* as the 3’ UTR of *Renilla* luciferase and a separate luciferase as a transfection control (**Fig. 1C**). We co-transfected HEK293 cells with plasmids expressing the dual luciferase reporters, Cas13b and gRNA expression cassettes or controls, and measured luciferase activity 24 hours later (**Fig. 1C-D**). All 15 gRNAs significantly silenced luciferase-DUX4 expression compared to the Psicheck2_DUX4 control (45-90.5% reduction; *P* < 0.0001, ANOVA) with gRNA1 (14.61 ± 1.4%), gRNA2 (13.64 ± 0.77%), gRNA6 (28.42 ± 1.76%), gRNA7 (31.44 ± 2.57%), gRNA8 (28.1 ± 0.81%), gRNA10 (20.89 ± 0.35%), gRNA14 (9.52 ± 1.81%), and gRNA15 (25.18 ± 3%) producing the most efficient luciferase silencing *in vitro* (**Fig 1D**). We did not observe luciferase-DUX4 silencing in HEK293 cells transfected with the dual-luciferase reporter and Cas13b alone or Cas13b with a control gRNA (nontar gRNA), which lacks sequence homology to *DUX4* and instead targets the *Gaussia* luciferase (Gluc) open reading frame (*62*). These luciferase results and controls confirmed specificity of our *DUX4*-targeting gRNAs (**Fig 1D**). For a secondary screen, we validated the silencing efficacy of Cas13b-gRNA against the wild-type *DUX4* target, employing western blotting to measure DUX4 protein levels (**Fig 1E-F**). To achieve this, we again performed co-transfection experiments but replaced Psicheck2_DUX4 with an expression plasmid containing a CMV promoter driving a V5 epitope-tagged, full-length *DUX4* construct. We extracted total proteins 20 hours later and performed western blots probed with antibodies to V5 (DUX4) or α-tubulin (loading control) (**Fig 1E-F**). We detected significant *DUX4* silencing in response to 13 of 15 guide RNAs, ranging from 40.93 ± 8.5% to 86.67 ± 3.6% reduction in DUX4 protein (**Fig 1E-F**) (P<0.05 to P < 0.0001, ANOVA, N = 3), confirming the findings from the luciferase assay. Specifically, compared to the nontar control, DUX4 protein levels were present at reduced levels in cells treated with gRNA1 (13.33 ± 3% remaining DUX4 protein), gRNA2 (27 ± 7.9%), gRNA6 (27.3 ± 15.58%), gRNA7 (26 ± 3.72%), gRNA8 (29.8 ± 9.8%), gRNA9 (45.26 ± 5.36%) gRNA14 (28.66 ± 6.04%), and gRNA15 (25.33 ± 6.6%) (**Fig 1E-F)**. As additional outcomes to support our western blot results, we also assessed the impacts of Cas13b with gRNA 1 and gRNA2 on DUX4 protein expression using a DUX4-GFP fusion protein construct and immunofluorescence (anti-DUX4-V5 antibodies) (**Fig 1G-H**). These experiments demonstrated cell-to-cell consistency of DUX4 knockdown at the protein level and confirmed our western blot data (**Fig 1E-F)**.

As a final efficacy screen in HEK293 cells, we assessed the ability of the CRISPR-Cas13b system to protect HEK293 cells from DUX4-induced cell death, using cell viability (trypan blue cell counts) and apoptosis (caspase-3/7) assays. While 8 of 15 gRNAs significantly improved cell viability compared to controls (gRNAs 1, 2, 3, 6, 8, 9, and 14) (**Fig 1I**), only 4 provided significant protection from apoptosis as determined by reduced caspase-3/7 activity (gRNAs 1, 2, 6, and 9) (**Fig 1J**). Taken together, our *in vitro* screens demonstrated that gRNAs 1 and 2 provided the most robust and consistent DUX4 knockdown, accompanied by protection from DUX4-induced cell death (**Fig 1**). We therefore chose gRNA1 and gRNA2 as our lead sequences for subsequent investigations.

### In vitro efficacy testing of lead guide RNAs in FSHD patient-derived muscle cells

Translating CRISPR-Cas13b as a FSHD therapy will ultimately require targeting human muscle cells, and we therefore next assessed the impact of *DUX4* knockdown by Cas13b in DUX4-expressing human myoblasts. Although *DUX4* expression should be the most direct indicator of target engagement for potential drug or gene therapy candidates, it is rare, transitory, and difficult to detect in FSHD muscle biopsies. FSHD patient-derived muscle cells also show a similar phenotype, in which *DUX4* is absent in most FSHD myoblasts at any given time (*23, 72, 73*). As an example relevant to this study, we used 15A myoblasts harvested from an FSHD patient muscle biopsy (gift of Dr. Charles Emerson, University of Massachusetts) (*72*). Although all 15A myonuclei are genetically capable of expressing *DUX4*, empirically only a small percentage produce *DUX4* at any given time (<0.1% to ∼3%) (*23, 51, 72*). These observations in FSHD biopsies and cell lines support that direct measurement of *DUX4* expression in human muscle biopsies is currently an unreliable outcome measure for FSHD clinical trials. As an alternative, the FSHD field has turned to measuring known DUX4-activated target genes as potential disease and therapy biomarkers (*27, 74-79*). We selected three established DUX4 target genes and FSHD disease biomarkers—*TRIM43*, *MBD3L2*, and *PRAMEF12*—based on their consistent differential expression between FSHD and healthy control cells in previous studies, and their natural expression in 15A myoblasts, which increases with differentiation into myotubes (*23, 48, 51, 74, 75, 80*). We therefore next evaluated the efficacy of our lead Cas13b gRNAs to suppress these 3 transcripts in 15A myoblasts (**Fig 2A**). To achieve this, we co-transfected 15A myoblasts with plasmids expressing Cas13b and gRNA1, gRNA2, or a nontargeting gRNA control (nontar), and induced myotube differentiation for 7 days. We then collected RNA and measured levels of *TRIM43*, *MBD3L2*, and *PRAMEF12* using quantitative RT-PCR. Compared to cells treated with Cas13b and the nontar-gRNA, both gRNA1 and gRNA2 significantly reduced levels of each biomarker by at least 76.43% (gRNA2) or 84.26% (gRNA1) (P < 0.0001, N=3 or N=4 independent experiments) (**Fig 2A**).

**Fig 2.**
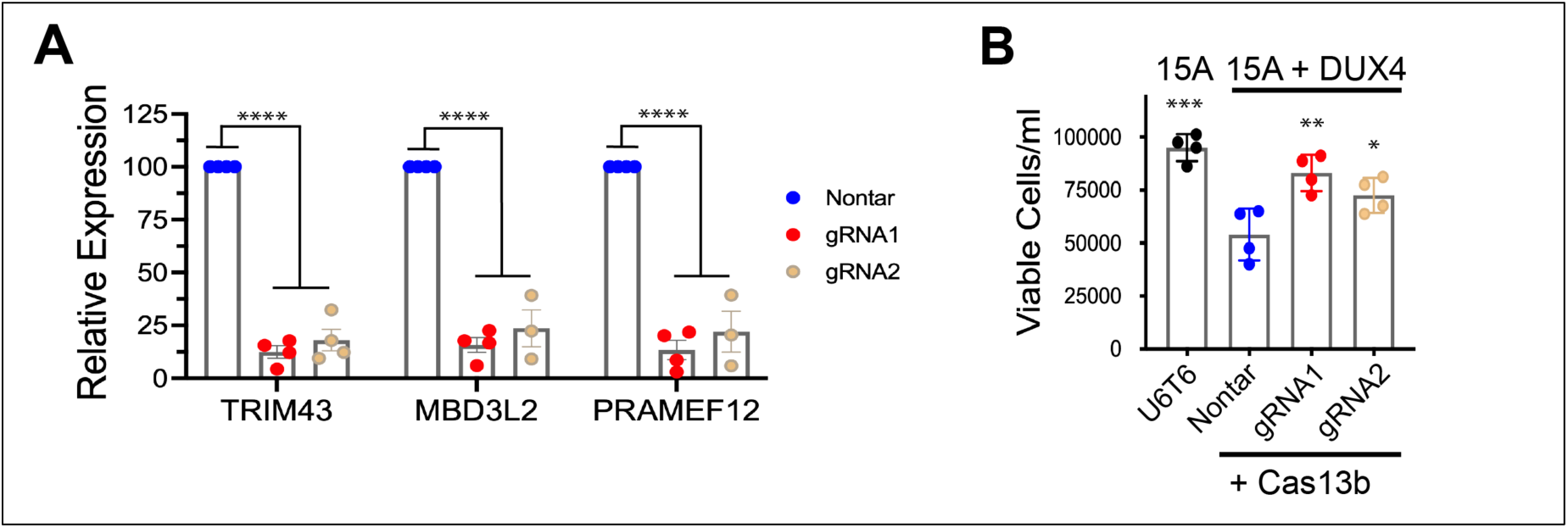
Lead gRNAs protect against cell death in human myoblasts and reduce the DUX4-activated biomarkers. (A) 15A FSHD patient-derived myoblasts were transfected with plasmids expressing wild-type Cas13b and each gRNA, and differentiated into myotubes. Quantitative RT-PCR was performed for DUX4-activated biomarkers *TRIM43*, *MBD3L2*, and *PRAMEF12*. Gene expression was normalized to *RPL13A*. While both gRNA 1 and gRNA2 significantly reduced levels of DUX4-activated biomarkers, all biomarkers were slightly but not significantly lower in cells treated with gRNA1 (12.46 ± 2.95% for *TRIM43*, 15.73 ± 3.48% for *MBD3L2*, and 13.39 ± 4.54% for *PRAMEF12*) vs gRNA2 (numbers?). (B) Cell viability significantly increased in gRNA1- and gRNA2-treated 15A myoblasts compared to nontar-gRNA controls, assessed 48 hours post-transfection via trypan blue counting assay. For qRT-PCR and cell viability, n=3 or 4 independent experiments were performed in triplicate as shown in each dot plot graph. *P < 0.05, **P < 0.01, ***P < 0.001, and ****P < 0.0001, ANOVA. U6T6: empty plasmid carrying the U6 promoter with no miRNA.

As a second outcome measure using patient-derived muscle cells, we assessed the ability of the CRISPR-Cas13 system to protect 15A myoblasts from DUX4-induced cell death. Because the majority of 15A myoblasts lack DUX4 expression, we used a co-transfection strategy to generate a more robust model system in which DUX4 was expressed more broadly and at levels sufficiently high to produce a cell death outcome. To do this, we co-transfected 15A human FSHD myoblasts with plasmids expressing *DUX4*, Cas13b, and our lead (gRNA1 and gRNA2) or control (nontar) gRNAs, and then assessed cell viability 48 hours later by counting living, trypan blue-negative cells (**Fig 2C**). *DUX4* transfection without Cas13b/gRNA silencing significantly reduced the percentage of viable 15A cells by 56.9 ± 2% (compare U6T6 to nontar; p=0.001, ANOVA, N=4 experiments). In contrast, Cas13b transfected with either gRNA1 or gRNA2 significantly protected myoblasts from DUX4-induced death, although to different degrees, with gRNA1 (87.5± 1.3% viable cells, p= 0.002, ANOVA) performing better than gRNA2 (76.31± 1% viable cells, p=0.03, ANOVA) (**Fig 2B**).

### Off-target assessments in vitro: Cas13b lacked collateral cleavage activity but activated immune response-associated transcripts in human cells

Ideally, CRISPR-based gene therapy systems should be programmed to degrade only RNA from the target disease gene while avoiding destruction of non-target RNAs. However, on-target specificity is not guaranteed. Theoretically, Cas13 could degrade non-target RNAs if the guide RNA has complementarity to an unintended transcript (gRNA sequence-dependent off-target activity) or through non-specific destruction of cellular RNAs that may be in the vicinity of a programmed Cas13 enzyme (gRNA sequence-independent collateral activity). Several studies have reported robust collateral activity from LwaCas13a (*81-83*) and RfxCas13d (*83-85*) in eukaryotic cells, while others reported none (*61, 62, 86*). A recent study demonstrating collateral activity from PspCas13b spurred us to investigate this potential effect using the Cas13b system *in vitro* (*83*). To do this, we constructed a dual reporter plasmid containing destabilized GFP (dGFP) and destabilized mCherry (dmCherry) genes under the control of CMV and SV40 promoters, respectively (**Fig 3A**) (*84*). Since both reporter transcripts arise from the same plasmid, and are therefore in the same vicinity within the host cell, we reasoned that collateral activity would be detectable by diminution of fluorescence of the reciprocal, non-targeted fluorophore, similar to a previously published study (*84*). To assess the potential for Cas13b-mediated collateral activity, we co-transfected HEK293 cells with the dual reporter plasmid and Cas13b, along with gRNAs specific to either GFP or mCherry, or a non-targeting gRNA control. Twenty-four hours later, the relevant target fluorophores were significantly reduced by Cas13b programmed with gRNAs complementary to GFP or mCherry, but we did not detect collateral reduction in the reciprocal reporter signal (**Fig 3A**).

**Fig 3.**
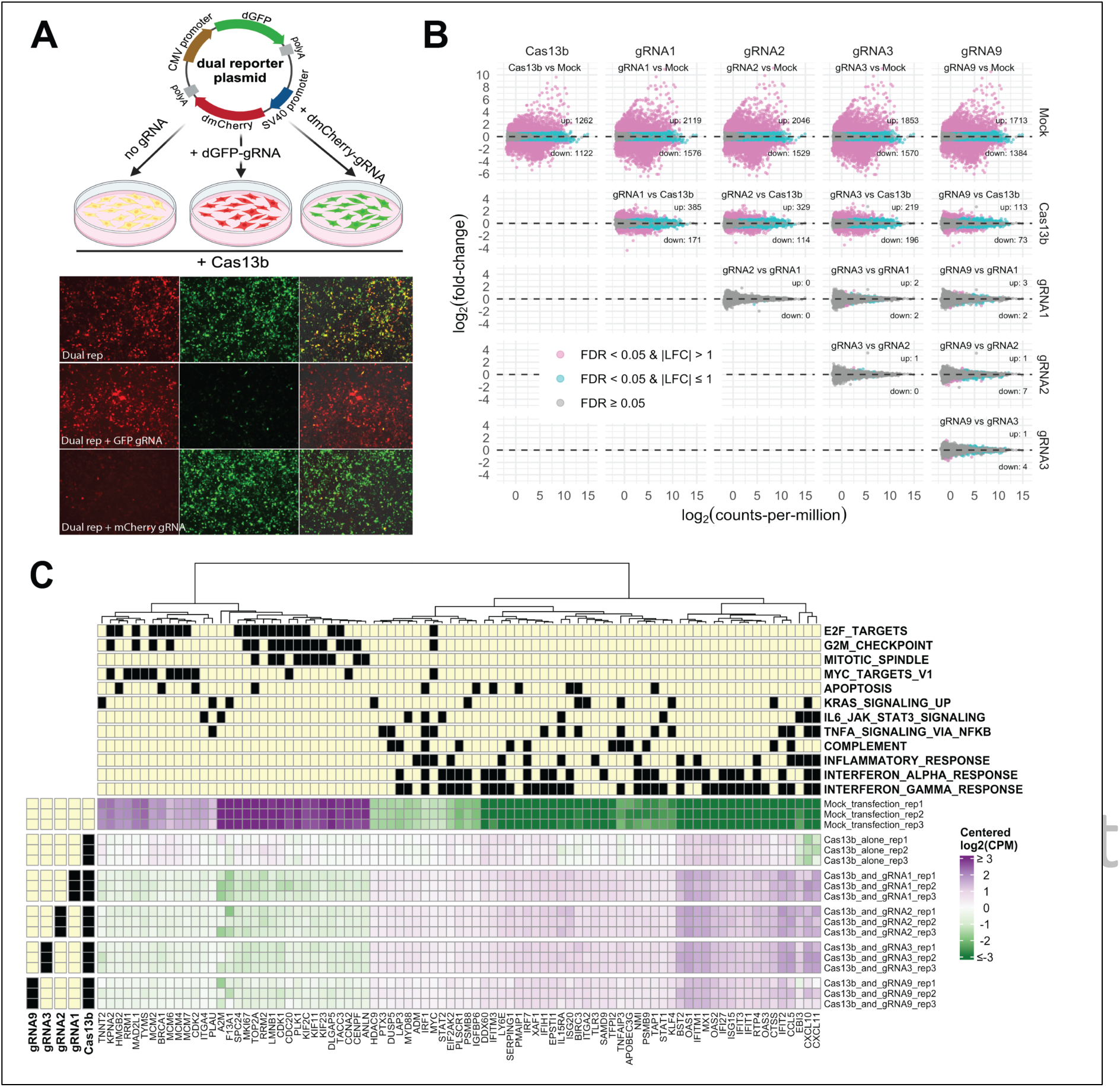
Collateral activity and off-target analysis of CRISPR-Cas13b system. (A) No collateral activity of Cas13b was detected in HEK293 cells using a dual reporter plasmid containing destabilized GFP (dGFP) and destabilized mCherry (dmCherry) genes under the control of CMV and SV40 promoters, respectively. 10X objective. (B) RNA sequencing to detect off-target effects of Cas13b with or without gRNAs in FSHD myoblasts (15A) 48 hrs post-transfection. Mock-transfected 15A myoblasts were used as a control. These MA plots show average log2(counts-per-million) (average expression across all samples) vs log2(fold-change) (or LFC for short) for 15 comparisons between pairs of conditions. Each point represents a single gene (19,834 total), and genes are colored magenta if test of differential expression had FDR < 0.05 as well as abs(LFC) > 1; cyan if test had FDR < 0.05 but not abs(LFC) >1; and grey otherwise. The number of genes colored magenta are labelled in each panel of the plot, separately for the up (LFC > 0) and down (LFC < 0) directions. As this cutoff there were 2,384 DEGs for the comparison of Cas13b alone vs Mock;, 3,097-3,695 DEGs for Cas13b with any gRNA vs Mock; 186-556 DEGs for Cas13b with any gRNA vs Cas13b alone; 0-8 DEGs for any gRNA vs another gRNA (all with Cas13b). (C) RNA-seq heatmap. This heatmap shows expression values in each sample for select genes in each of the HALLMARK gene sets that were significantly enriched (p < 10-4) for DEGs in one or more of the between-group comparisons. These sets are indicated along the top margins of the heatmap, where a black square denotes that the gene labeled in the bottom margin belongs to the gene set. As these gene sets can have up to ∼200 genes and up to ∼100 DEGs, only a subset of genes from the sets are shown: the 10 genes per gene set with the smallest p-value in any of the between-group comparisons, just among genes with abs(LFC) > 1. There can be more than 10 displayed genes belonging to some of the gene sets because the HALLMARK gene sets can overlap each other, as evident from the grid in the upper margin. The grid in the left margin highlights whether samples included Cas13b alone, Ca13b together with a guide RNA, or neither (see also the sample names in the right margin). The displayed expression values are counts-per-million (CPM) on a log2 scale (using prior.count=2 in edgeR to avoid log of zero) that are centered to have mean of zero for each gene; thus, as shown in the color-bar to the right, purple denotes expression that is higher in a sample than the average across all samples, and green denotes expression that is lower in a sample than the average across all samples. The color-scale for centered log2(CPM) is capped at values of +/-3, corresponding to 8-fold above and below the average across all samples, since without a cap the wide dynamic range would make differences of +/-1 (i.e., 2-fold differences) difficult to discern.

Next, to assess potential non-specific activity of Cas13b on global transcripts in FSHD patient cells, we transfected 15A myoblasts with Cas13b alone or with lead gRNAs (gRNA1 and gRNA2, which both bind sequences encoded by exon 1 of *DUX4* and overlap each other by 22 nt). We also used two other guide RNAs: gRNA3, which also binds exon 1 of *DUX4* and overlaps gRNA2 by 4 nt; and gRNA9, which binds exon 2 of *DUX4* and is located distant from gRNA1-gRNA3 binding sites in both the primary sequence and predicted secondary structure at the binding site of the *DUX4* mRNA (**Fig 1**). In addition, we performed a mock transfection as an additional control, for a total of six experimental conditions each with three replicates. After 48 hours, we harvested RNA, and prepared libraries for whole transcriptome RNA sequencing at a depth of ∼20 million paired-end reads per sample. We performed tests of differential gene expression between all pairs of conditions. Using cutoffs of false-discovery rate (FDR) < 0.05 and abs(fold-change) > 1 for differentially expressed genes (DEGs), we found widespread differences between the mock transfection and all other conditions (2,384 DEGs for mock vs Cas13b alone and 3,097-3,695 DEGs for mock vs Cas13b with any guide RNA). In addition, there were intermediate levels of differences between Cas13b alone and Cas13b with any guide RNA (186-556 DEGs); and only mild differences between guide RNAs (0-8 DEGs) (**Fig 3B, Table1, Table S2**).

**Table 1.** RNA-seq differential expression summary. This table lists the number of differentially expressed genes (DEGs) for each of 15 between-group contrasts at 3 levels of stringency: **(DE_1)** FDR < 0.05 for glmQLFTest; **(DE_2)** FDR < 0.05 and abs(LFC)>1 for glmQLFTest; **(DE_3)** FDR < 0.05 for glmTreat with option lfc=1. Counts are given separately for genes with LFC > 0 (direction = up) and LFC < 0 (direction = down).

| **Contrast** | **DE_1** | **DE_1** | **DE_2** | **DE_2** | **DE_3** | **DE_3** |
| --- | --- | --- | --- | --- | --- | --- |
| **Direction** | **up** | **down** | **up** | **down** | **up** | **down** |
| Cas13b_vs_Mock | 4568 | 4880 | 1262 | 1122 | 538 | 630 |
| gRNA1_vs_Mock | 5900 | 5435 | 2119 | 1576 | 971 | 1012 |
| gRNA2_vs_Mock | 5865 | 5419 | 2046 | 1529 | 989 | 998 |
| gRNA3_vs_Mock | 5597 | 5392 | 1853 | 1570 | 933 | 1013 |
| gRNA9_vs_Mock | 5305 | 5246 | 1713 | 1384 | 802 | 895 |
| gRNA1_vs_Cas13b | 2261 | 2597 | 385 | 171 | 15 | 14 |
| gRNA2_vs_Cas13b | 2104 | 2387 | 329 | 114 | 16 | 12 |
| gRNA3_vs_Cas13b | 1775 | 2182 | 219 | 196 | 20 | 23 |
| gRNA9_vs_Cas13b | 1107 | 1398 | 113 | 73 | 0 | 0 |
| gRNA2_vs_gRNA1 | 0 | 2 | 0 | 0 | 0 | 0 |
| gRNA3_vs_gRNA1 | 17 | 25 | 2 | 2 | 0 | 0 |
| gRNA9_vs_gRNA1 | 19 | 34 | 3 | 2 | 0 | 0 |
| gRNA3_vs_gRNA2 | 14 | 8 | 1 | 0 | 0 | 0 |
| gRNA9_vs_gRNA2 | 45 | 96 | 1 | 7 | 0 | 0 |
| gRNA9_vs_gRNA3 | 51 | 85 | 1 | 4 | 0 | 0 |

We performed gene set analysis on the DEGs, comparing them to the MSigDB collections of genes sets. Using a cutoff of P-value < 10^-4^ for over-representation (see RNA-seq methods), the HALLMARK gene sets enriched for DEGs *up* in the Cas13b vs mock comparison were INTERFERON_GAMMA_RESPONSE, INTERFERON_ALPHA_RESPONSE, COMPLEMENT, INFLAMMATORY_RESPONSE, and TNFA_SIGNALING_VIA_NFKB, suggesting an immune or inflammatory response to transfection with Cas13b (**Fig 3C; Table S3**). The HALLMARK gene sets enriched for DEGs *down* in the Cas13b vs mock comparison were E2F_TARGETS, G2M_CHECKPOINT, and MITOTIC_SPINDLE, suggesting disruptions to cell-cycle in response to transfection with Cas13b (**Fig 3C; Table S3**). For all four guide RNAs, the same HALLMARK gene sets showed the strongest enrichment in the Cas13b with gRNA vs mock comparisons as in the Cas13b alone RNA vs mock comparisons (**Fig 3C; Table S3**). These findings may not be surprising since these comparisons are all relative to the same mock samples, with many DEGs in common (952 upregulated and 935 downregulated DEGs shared by all five comparisons of Cas13b alone or with a guide RNA vs mock). One additional HALLMARK gene set, MYC_TARGETS_V1, was also enriched (P-value < 10^-4^) among the downregulated DEGs for gRNA1 and gRNA2 (and had P-value < 0.01 for the other two guide RNA) but not for Cas13b alone (P-value = 0.11).

In direct comparisons of Cas13b with guide RNAs to Cas13b alone, INTERFERON_GAMMA_RESPONSE, INTERFERON_ALPHA_RESPONSE, COMPLEMENT and INFLAMMATORY_RESPONSE were among the HALLMARK gene sets most enriched for DEGs that had higher expression with a guide RNA added than with Cas13b alone (although in some cases not with P-value < 10^-4^), and E2F_TARGETS and G2M_CHECKPOINT were significantly enriched for DEGs that had lower expression with any guide RNA added than with Cas13b alone. This suggests that the alterations related to immune and inflammatory responses and cell-cycle seen in the comparisons between Cas13b alone and mock transfection may be exacerbated by addition of a guide RNA. One additional gene set, IL6_JAK_STAT3_SIGNALING, was enriched for DEGs that had higher expression with any guide RNA added than with Cas13b alone, and KRAS_SIGNALING_UP and APOPTOSIS were significant for some guide RNA and near-significant for some others. No HALLMARK gene sets were significantly enriched for DEGs from comparisons between guide RNAs, which is unsurprising given the small number of DEGs in these comparisons.

Cells in these experiments had very low levels of *DUX4* and well-characterized markers of DUX4 activity such as the *TRIM43*, *MBD3L2*, and *PRAMEF12,* which we measured in quantitative RT-PCR assays for screening guide RNAs. Typically, we found 0, and at most 4 read counts, for any of these genes in any of the mock or Cas13b-only samples (out of ∼10M-15M read counts total per sample), and levels were not significantly reduced in samples with guide RNAs added. These results were consistent with reports that myoblasts from individuals with FSHD express lower levels of *DUX4* and DUX4-induced-genes than myotubes (*27, 74, 87*). Thus, it seems unlikely that the DEGs in the Cas13-with-guide RNA versus Cas13b-alone comparisons are consequences of reducing levels of DUX4 expression or activity. It moreover seems unlikely that the DEGs primarily reflect sequence-specific off-target silencing of other genes since there are so few DEGs in comparisons between different guide RNAs, which should pick up sequence-specific off-target effects unless both guide RNAs being compared happen to target the same gene. While this could frequently occur for gRNA1 and gRNA2, which overlap by 22 nt, gRNA3 has little overlap with gRNA1 and 2 (sharing just 4 nt with gRNA2 and none with gRNA1) and gRNA9 has no overlap at all with the others. Aside from *DUX4* itself there are only two genes that have more than 10 nt of complementarity with all four guide RNAs, *AC003681.1* and *ADGRV1,* both of which have exactly 11 nt of complementarity with each of the four gRNAs. While the expression levels of *AC003681.*1 did not differ much between conditions, the expression for *ADGRV1* was reduced between 1.7- and 2.8-fold in the comparisons of Cas13b with guide RNA versus Cas13b alone; it is not clear whether this could plausibly be directly due to off-target binding, or – even if so – how much this might contribute to the hundreds of DEGs in those comparisons. Even in comparisons between different guide RNAs, which had few DEGs, the DEGs did not appear to be directly caused by off-target binding, as none of the DEGs had more than 9 nt of complementarity with either of the guide RNAs being compared.

### Cas13b-gRNA prevents *DUX4*-mediated muscle damage *in vivo*

Having demonstrated that gRNA-programmed Cas13b could reduce *DUX4* mRNA and protein levels, and prevent several DUX4-associated outcomes *in vitro*, we next tested the system *in vivo*. To do this, we generated three AAV6 vectors: two expressing gRNAs (gRNA1 or a non-targeting control) under a human U6 promoter (AAV6.U6-gRNA1 or AAV6.U6-nontar), and a third carrying a PspCas13b open reading frame (ORF) driven by a minimal CMV (miniCMV) promoter (AAV6.miniCMV-Cas13b) (**Fig 4A**). In addition, we used our previously published AAV-based *DUX4* mouse model for initial *in vivo* efficacy testing and therefore made a fourth AAV6 vector expressing a V5 epitope-tagged *DUX4* ORF and 3’ UTR (*89*). To test the potential protective effects of the gRNA1-programmed Cas13b system *in vivo*, we next co-delivered *DUX4*, gRNA1 and Cas13b expression vectors into wild-type C57BL/6 tibialis anterior (TA) muscles via intramuscular injection. Three weeks after injection, muscles treated with *DUX4*, Cas13b and the non-targeting gRNA showed widespread histopathology, including mononuclear cell infiltration and abundant centrally nucleated (CN) myofibers, indicating muscle damage and repair (**Fig 4B**). Conversely, gRNA1-treated muscles were protected from similar histopathology, indicated quantitatively by significant reductions in myofibers containing central nuclei, demonstrating that Cas13b can prevent DUX4-induced muscle damage *in vivo* (4.6-fold reduction in central nuclei, P<0.001; t-test) (**Fig 4B-C**). We also measured decreased *DUX4* mRNA expression in Cas13b/gRNA1-treated muscles compared to controls using ddPCR (1.85-fold reduction, P<0.05, N=6) and RNAscope (41.15-fold reduction, P<0.001, N=9) (**Figs 4B,D,E**). These differences were associated with significant reductions in the DUX4-activated mouse biomarker *Wfdc3* (2.9-fold reduction, P<0.0001, N=8) (**Fig 4F**). Additionally, we detected a reduction in DUX4-V5 and cleaved caspase-3 signals in treated muscles via immunofluorescent staining (**Fig S2**). We also confirmed the presence of gRNA1 and Cas13b mRNA expression, measured by real-time PCR two weeks after injection (**Fig 4G-H**). These results support that Cas13b can potently silence *DUX4* mRNA and suppress DUX4-associated pathology *in vivo*.

**Fig 4.**
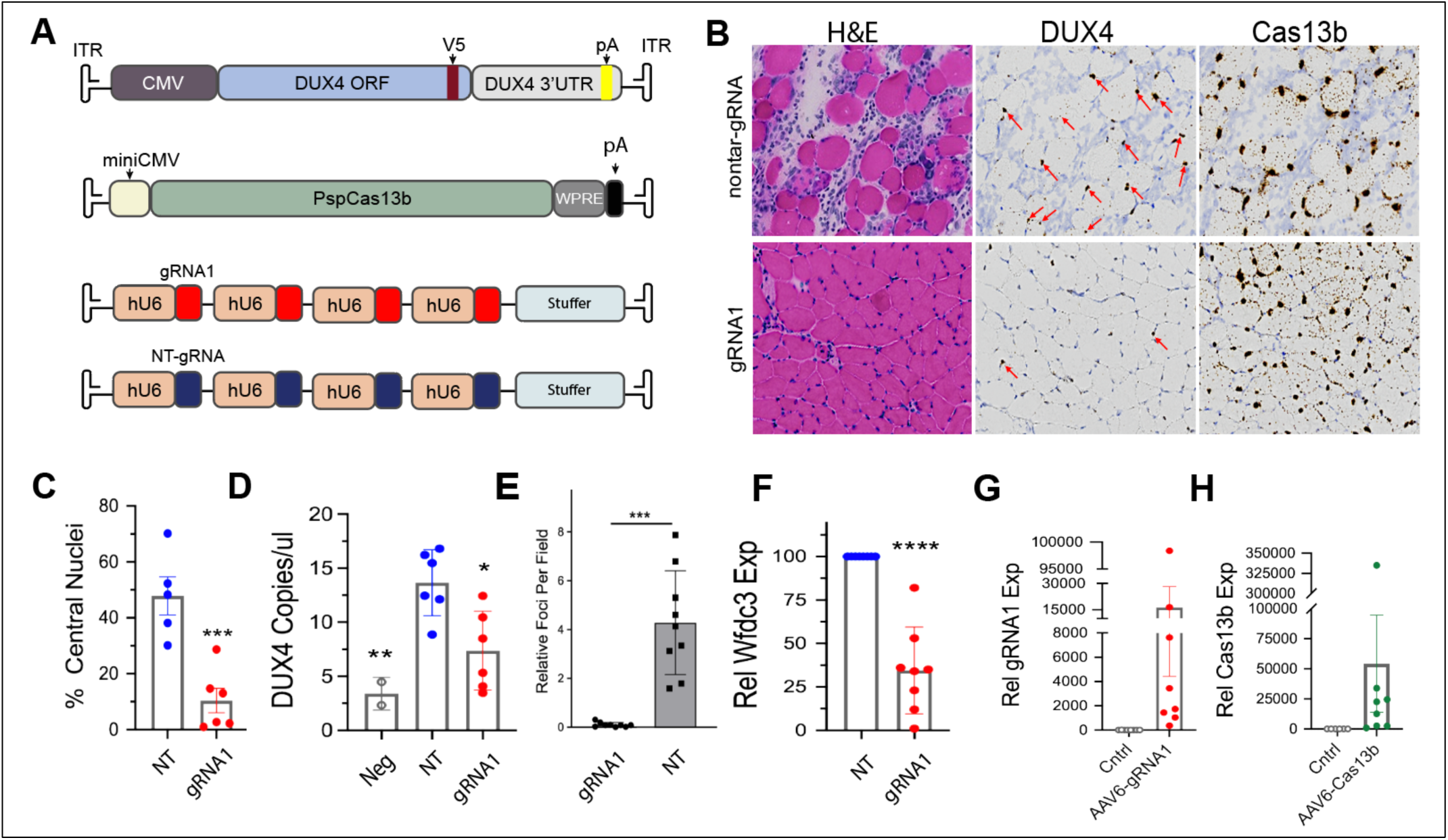
CRISPR-Cs13b improved DUX4-associated pathology in FSHD mice. (A) Schematic drawing of AAV expression cassettes carrying *DUX4*, Cas13b, or gRNA genes. (B-E) By 3 weeks, treated muscles demonstrated reduced histopathology and degeneration/regeneration compared to untreated muscles. Arrows indicate nuclei positively stained for *DUX4* or Cas13b mRNA using RNAscope *in situ* hybridization assay. 20X objective. (C) Cas13b with gRNA1 produced a 4.6-fold reduction in the percentage of myofibers containing centrally located nuclei (CN); (D) a 1.85-fold reduction in *DUX4* expression (ddPCR), N=6, and (E) 41.2-fold reduction in *DUX4* mRNA foci stained by RNAscope N=9, in treated muscles. (F) Cas13b + gRNA1-treated muscles showed significantly reduced expression of *Wfdc3*, a DUX4-activated biomarker in mice (*Wfdc3*) (2.9-fold reduction, N=8). (G-H) qRT-PCR analysis demonstrated abundant gRNA1 (3,955-fold, N=8) and Cas13b (32,965-fold, N=8) expression in injected muscles. *P < 0.05, **P < 0.01, ***P < 0.001, and ****P < 0.0001. NT: nontar-gRNA, Neg: untreated negative control, Cntrl: untreated wild-type control.

### Loss of sustained AAV6.Cas13b expression is associated with immune response in mice

Our *in vivo* studies showed promising protection from DUX4-mediated pathology in mouse muscle over a short time point (3 weeks) in an aggressive FSHD model, but achieving sustained efficacy from a CRISPR-Cas13-based gene therapy for FSHD will require long-term expression of Cas13b and gRNAs *in vivo*. We therefore performed an experiment in wild-type mice to measure sustained expression of Cas13b and gRNA1 over time, thereby supporting translatability of our CRISPR-Cas13 gene therapy for FSHD. To do this, we IM-injected AAV6.Cas13b and AAV6.gRNA1 expression vectors into adult C57BL/6J TA muscles and harvested RNA 2, 4, 8, and 24 weeks (6-month) later. Using real-time PCR, we found that Cas13b and gRNA1 levels rose between 2- and 4-weeks, and again from 4- to 8-weeks after injection. However, at the 24-week (6-month) timepoint, both Cas13b (2.44-fold, P<0.0001, N=9) and gRNA1 (1.99-fold, P<0.01, N=9) expression were significantly reduced (**Fig 5A**). We also noted focal mononuclear cell infiltration in DUX4-negative C57BL/6J muscles injected solely with Cas13b, beginning 5 weeks post-injection (**Fig 5B**). These results suggested that potential host immune responses to Cas13b could be responsible for its reduction between 8- and 24-weeks. To investigate this possibility, we first stained wild-type TA muscle sections with fluorophore-labeled anti-T cell antibodies, 5 weeks after delivery of AAV6.Cas13b. We found CD3-, CD4-, and CD8-positive staining co-localizing with mononuclear infiltrates in H&E-stained sections (**Fig 5B**). To validate the immunofluorescence results, we again injected wild-type C57BL6 muscles with saline, AAV6.Cas13b, or AAV6.mi405, which expresses a *DUX4*-targeting miRNA from an AAV genome but does not produce a protein *(23, 52)*, and then isolated immune cells from muscle for flow cytometry. We detected increased percentages of muscle-infiltrating immune cells in AAV6.mi405- or AAV6.Cas13b-injected muscles compared to saline controls. These immune infiltrates showed significantly increased staining for T-cells (Thy1+/CD3+). While both AAV6.mi405- and AAV6.Cas13b-injected mice showed similar significant increases in activated helper T cells (CD69+/CD4+) compared to saline-injected muscles, only the AAV.Cas13b-injected muscles showed a significant increase in CD8+ T cells. These results suggested that activation of a cytotoxic T cell response directed at Cas13b-transduced muscle cells compromised the long-term expression of CRISPR components in our model system.

**Fig 5.**
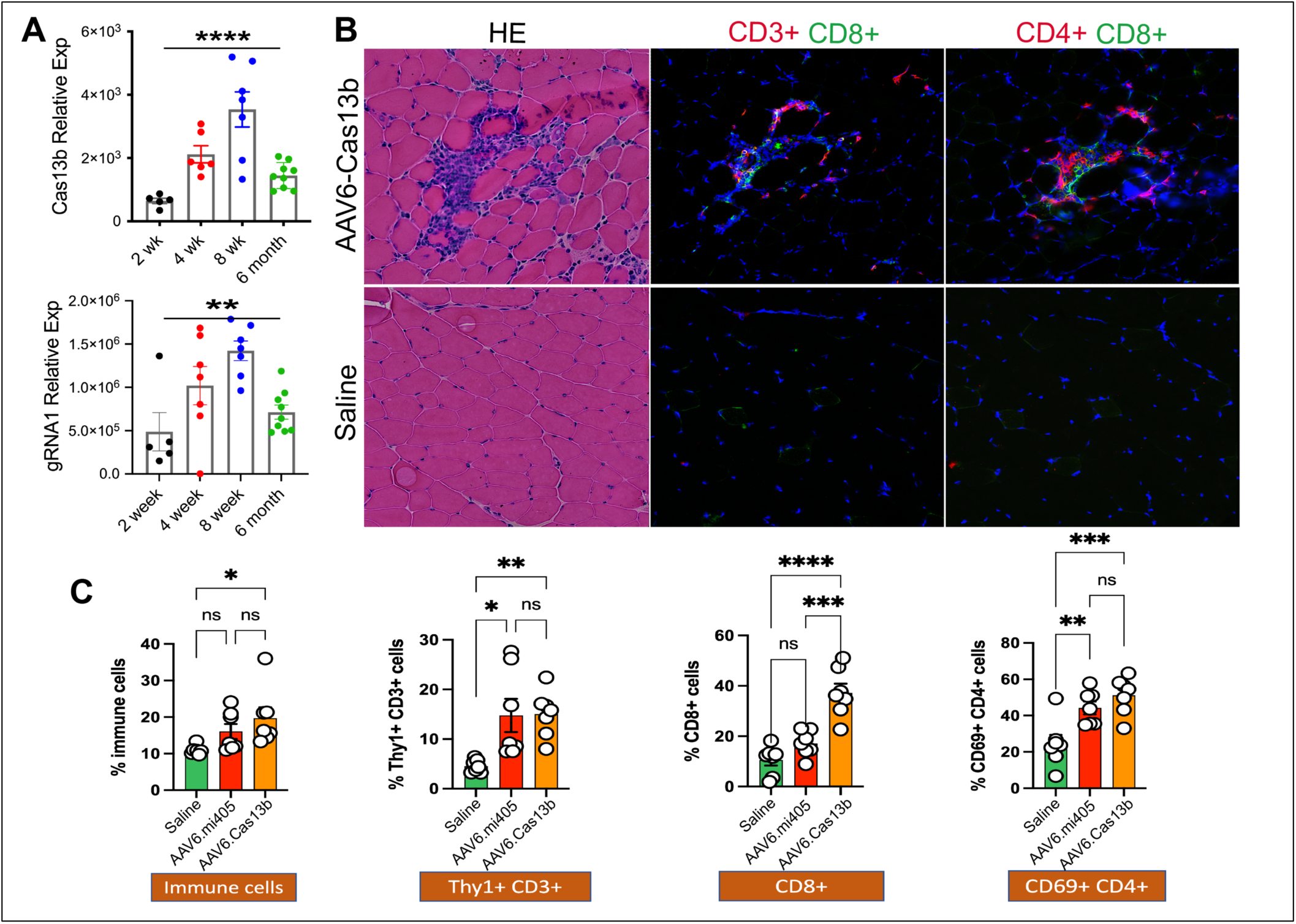
Immune response against AAV6-Cas13b in mice. (A) Cas13b and gRNA1 expression levels over a 6-month time span in adult C57BL/6J TA muscles following AAV6.Cas13b or AAV6.gRNA1 treatment. While Cas13b and gRNA1 expression levels continued to increase until 8 weeks post-injection, a significant reduction in each transcript level was detected at a 6-month time point (2.44-fold reduction of Cas13b and 1.99-fold reduction of gRNA between 8 wks and 6 months. N=5-9 per group) (B) Focal mononuclear cell infiltration was detected 5 weeks after injection of muscles with AAV6.Cas13b. CD3+, CD4+, and CD8+ cells co-localized with mononuclear infiltrates in H&E-stained sections, detected by fluorescent staining. (C) Flow cytometry analysis detected a significant increase in percentages of muscle-infiltrating immune cells in AAV6-Cas13b compared to saline-injected muscles. While both AAV6.mi405 (non-protein coding AAV) and AAV6.Cas13b injected muscles showed a significant increase in Thy1/ CD3+ and CD69+/CD4+ cells compared to saline-treated muscles, only AAV6.Cas13b-injected muscles showed significantly increased CD8+ T-cells. Together, these data suggest activation of a cytotoxic T-cell response to intramuscular AAV6-Cas13b injections in mice. N=7 per group. *P < 0.05, **P < 0.01, ***P < 0.001, and ****P < 0.0001, ANOVA (followed by Tukey’s post hoc tests for flow cytometry data).

## DISCUSSION

Translational studies on FSHD have historically lagged those of other major muscular dystrophies, such as DMD. Like DMD, the clinical features of FSHD were first described in the 1800s, and the disease was recognized as one of three major forms of muscular dystrophy in the pre-molecular biology days of 1954 (*90, 91*). The search for both the DMD and FSHD genes began in earnest in the 1980s, with foundational positional cloning studies leading to the 1986 discovery of the DMD gene, and identification of the *mdx* mouse, which provided a critical tool for testing countless therapies (*91*). These milestones ultimately enabled several approved treatments for DMD, although none yet represent complete “cures” of the disease, and improvements are indicated. In contrast, the 1992 linkage of FSHD to a deletion on human chromosome 4q35 failed to identify an obvious FSHD gene (*91*), and unfortunately, the pathogenic mechanisms underlying FSHD required nearly 2 more decades of study before the *DUX4* gene was recognized as a primary insult. The lack of understanding of FSHD pathogenesis negatively impacted model and therapy development. Importantly however, during the last ∼15 years the FSHD field has been making up for lost time. We now have better understanding of FSHD molecular mechanisms leading to *DUX4* de-repression, with several animal and cell models now available (*13, 15, 19, 25-28, 30, 32, 34, 37-40, 42*), robust clinical trial readiness studies ongoing (*92, 93*), and numerous FSHD therapies in pre-clinical development (*44-60*) or clinical trials (*94, 95*) (NCT05397470, NCT04003974, NCT02603562, NCT02239224).

Current therapeutic strategies for FSHD are largely, but not exclusively, focused on inhibiting *DUX4* gene expression or interfering with the activity of the DUX4 protein. At the DNA level, *DUX4* inhibition strategies include approaches to epigenetically silence *DUX4* by increasing D4Z4 DNA methylation using CRISPRi (*44, 46*) or disabling the FSHD-permissive *DUX4* poly(A) signal using CRISPR/Cas9-mediated DNA editing (*45, 67, 68*). Another recent approach involved testing a small molecule inhibitor of p38 MAP kinase called Losmapimod, which blocks *DUX4* expression (*96*) *in vitro* and *in vivo* (*95, 97*) but failed to achieve FDA approval after a Phase III clinical trial. Numerous ongoing strategies focus on inhibiting *DUX4* messenger RNA using RNA interference (RNAi), poly(A) suppression, or RNAseH-mediated gene silencing, including various oligonucleotide-based approaches *(23, 47-57)* and, in our laboratory, gene therapy using an engineered miRNA called mi405 (*52, 89*). In the study described here, we tested an alternative approach to suppress *DUX4* mRNA using CRISPR/Cas13 gene therapy, because some previous reports suggested CRISPR/Cas13 is a superior silencing system to RNAi (*61, 62, 86, 98*). We found that CRISPR/Cas13 was indeed capable of directing robust silencing of *DUX4*, and offered protection from some myopathic phenotypes in DUX4-expressing mice 3 weeks after injection. Unfortunately, this protective effect was not sustained, as we noticed histological muscle lesions in AAV6.Cas13b-injected mice at later time points (**Fig 5**). In contrast, we have never observed such a loss of efficacy over time following delivery of AAV.mi405 to hundreds of mice using various AAV capsids. We hypothesized that this lack of sustained protection could arise from loss of Cas13b expression mediated by a host immune response (*99*). We therefore investigated this possibility in wild-type mice lacking the complicating factor of toxic *DUX4* expression. Five weeks after injecting wild-type mouse muscles with AAV6.mi405 or AAV6.Cas13b, only Cas13b-injected muscles contained elevated CD8+ mononuclear cells, thereby suggesting a cytotoxic T lymphocyte (CTL) response to the Cas13b protein. Our *in vitro* RNA-seq off-target data supported the *in vivo* findings, as several immune response pathways were significantly increased in human myoblasts transfected with Cas13b expression constructs (**Fig 3C; Table S3**). Interestingly, using KEGG Legacy gene set analysis, we detected HLA-B, HLA-C, B2M, TAP1, HLA-F, HLA-A, HLA-E, and TAP2 DEGs (Table S3) which suggests that antigen presentation is induced in muscle, reinforcing the interpretation that CD8+ cytotoxic T lymphocytes (CTLs) are reacting to muscle cells presenting Cas13 peptides. Together, these findings would represent the first to suggest the possibility of immune-mediated clearance of an AAV-delivered Cas13b gene therapy product *in vivo*, but our data are not the first to suggest an immune response to a CRISPR system (*99-103*). Specifically, an AAV.Cas9-mediated gene editing study in DMD dogs showed robust dystrophin expression that was accompanied by muscle inflammation and CTL response (*101*). Consistent with our Cas13b study, this group also reported cytotoxic T cell responses against Cas9 in wild-type dogs, while other transgenes (micro-dystrophin, SERCA2a, alkaline phosphatase) failed to elicit such responses (*101*). These findings, coupled with a recent report that humans have pre-existing immunity to Cas13d derived from *Ruminococcus flavefaciens* bacteria (*99*), suggest that immune responses could impede development of CRISPR-based gene therapies for neuromuscular disease and perhaps any therapies involving *in vivo* delivery of CRISPR effectors. While *ex vivo* CRISPR editing for cell therapy shows promise, we propose that more *in vivo* studies in neuromuscular disease and are needed to better understand immunity to CRISPR proteins. In our opinion, until more confidence is gained to support the safety and lack of deleterious immunogenicity to Cas proteins, *in vivo* CRISPR-based therapies should proceed with caution to mitigate clearance of gene therapy products, or in a worst case, avoid potentially catastrophic immune responses in humans. Further optimization of the CRISPR system including development of engineered Cas proteins with reduced immunogenicity (*104-108*) and designing more specific gRNAs and Cas proteins (*109-117*) may offer improvements for future safe applications. Additionally, the use of tissue-specific promoters rather than ubiquitous viral promoters, such as CMV, may prevent expression of Cas proteins in antigen-presenting cells, and thereby reduce subsequent cytotoxicity or immune response against these proteins.

## Materials and Methods

### Designing DUX4 targeting Cas13b-gRNAs

We obtained a previously described Cas13b expression plasmid in which the EF1a promoter drives expression of Prevotella sp. P5–125 (PspCas13b) Cas13b (Addgene #103862) (*62*). We designed 30-nt gRNAs following previously described methods (*62, 63*). Briefly, computational folding predictions of the *DUX4* mRNA were performed using the RNAfold program (http://rna.tbi.univie.ac.at/cgi-bin/RNAWebSuite/RNAfold.cgi). We designed fifteen gRNAs (Table S1) targeting predicted unstructured target sites on the *DUX4* mRNA, or sites for which we previously showed regulation by designed artificial miRNAs (*52, 118*). We then synthesized gRNA expression cassettes using the human U6 promoter on a pUCIDT plasmid backbone (Integrated DNA Technologies, Coralville, Iowa). The non-targeting control gRNA sequence targeting *Gaussia* luciferase (Gluc) (GCAGGGTTTTCCCAGTCACGACGTTGTAAAA; Table S1) was adapted from a previous study (*62*).

### Cell culture

HEK293 cells were cultured in DMEM supplemented with 10% fetal bovine serum, 1% L-glutamine, and 1% penicillin/streptomycin at 37°C in 5% CO_2_. Affected immortalized human myoblasts derived from an individual with FSHD (15Abic; gift of Dr. Charles Emerson, University of Massachusetts Chan School of Medicine) were expanded in DMEM supplemented with 16% medium 199, 20% fetal bovine serum, 1% penicillin/streptomycin, 30 ng/mL zinc sulfate, 1.4 mg/mL vitamin B12, 55 ng/mL dexamethasone, 2.5 ng/mL human growth factor, 10 ng/mL fibroblast growth factor, and 20 mM HEPES (*72, 119*). Cells were maintained as myoblasts and differentiated prior to measurement of *DUX4* mRNA and DUX4-activated biomarkers by qRT-PCR and RNAscope. To produce myotubes, transfected myoblasts were switched to differentiation medium composed of a 4:1 ratio of DMEM: medium 199 supplemented with 15% KnockOut Serum (Thermo Fisher Scientific), 2 mM L-glutamine, and 1% antibiotics/antimycobiotics for up to 7 days before harvesting.

### Dual-luciferase assay

We conducted this assay following the protocol described in a previous publication (*52*) and using the dual-luciferase reporter assay system from Promega, with slight modifications. The psiCheck2 dual luciferase reporter plasmid (Promega) contains separate *Renilla* and luciferase genes. Downstream of the *Renilla* luciferase stop codon, we cloned full-length DUX4 (DUX4-FL) including its open reading frame, two small introns, and untranslated exons 2 and 3, the latter of which contains the FSHD permissive poly(A) signal. *DUX4* sequences thus served as the *Renilla* luciferase 3’ untranslated region (UTR). The Firefly luciferase expressed under separate promoter acts as a control. We pre-plated HEK293 cells (42,000 cells/well) 16 h before transfection in 96-well plates. We then co-transfected cells with the luciferase DUX4 reporter (called psiCh-DUX4) and individual gRNA expression plasmids along with plasmid expressing Cas13b using Lipofectamine 2000 (Invitrogen). The luciferase activity was measured 48 hours after transfection, and we quantified DUX4 gene silencing by measuring the Renilla to Firefly ratio for each gRNA.

### Viability assay

HEK293 cells (250,000 cells/well) were seeded in 24-well plates and co-transfected (Lipofectamine 2000, Invitrogen) with an expression plasmid (CMV.DUX4-FL) along with plasmids expressing each gRNA and Cas13b. The cells were trypsinized 48 h after transfection and collected in 1 mL of growth medium. Automated cell counting was performed using Countess cell counting chamber slides and confirmed with traditional cell counting using a hemocytometer and trypan blue staining. Three independent experiments were performed, and data reported as a mean of total cell number ± SEM per group.

To evaluate myoblast viability, we electroporated (Lonza, VVPD-1001) CMV-DUX4 and CRISPR/Cas13 expression plasmids into 15Abic myoblasts (500,000 cells per reaction), following the protocol outlined in a previous study (*51*). After 48 hours, cells were trypsinized and counted as above.

### Apoptosis assay

HEK293 cells (42,000 cells/well) were plated on a 96-well plate 16 h prior to transfection. The next morning, cells were co-transfected (Lipofectamine 2000, Invitrogen) with CMV.DUX4-FL and U6-gRNA and EF1a-Cas13b expression plasmids, or controls. Caspase 3/7 activity was measured using the Apo-ONE Homogeneous Caspase-3/7 Assay (Promega, Madison, WI) 48 h after transfection using a fluorescent microplate reader (Spectra Max M2, Molecular Devices, Sunnyvale, CA). Three individual assays were performed in triplicate, and data were averaged per experiment and reported as mean caspase activity ± SEM relative to our control assay, which was transfected with CMV.DUX4-FL only.

### Western blot assay

To evaluate the impacts of the CRISPR/Cas13b system on DUX4 protein expression, we co-transfected HEK293 cells with CMV.DUX4-FL, with and without a V5 epitope tag at the C-terminus, along with each gRNA and Cas13b expression plasmids using Lipofectamine 2000 (Invitrogen). Twenty hours later, total protein was extracted using RIPA buffer (50 mM Tris, 150 mM NaCl, 0.1% SDS, 0.5% sodium deoxycholate, and 1% Triton X-100) supplemented with protease inhibitors (Thermo Fisher). Total protein concentration was determined using the DC Protein Assay Kit (Bio-Rad). Subsequently, 25 μg of each protein sample underwent electrophoresis on a 12% SDS-polyacrylamide gel followed by semi-dry transfer onto polyvinylidene fluoride (PVDF) membranes. Membranes were then blocked in 5% non-fat milk and incubated overnight at 4°C with primary monoclonal mouse anti-DUX4 (1:500; P4H2, Novus Biologicals), rabbit polyclonal anti-V5 (1:1000, ab9116, Abcam), or rabbit polyclonal anti-α-tubulin antibodies (1:1000, ab15246, Abcam). Following multiple washes, the blots were probed with horseradish peroxidase (HRP)-conjugated secondary antibodies (Jackson ImmunoResearch) for 1-2 hours at room temperature. Protein bands were visualized using X-ray film after incubation in Immobilon chemiluminescent HRP substrate (Millipore). Quantification of protein levels was performed using ImageJ software (National Institutes of Health, https://imagej.nih.gov/ij/).

### Immunocytochemistry of DUX4-V5

Immunocytochemistry staining was employed to validate the findings from the western blot assay. In this experiment, V5 epitope-tagged DUX4 protein was visualized in transfected cells using immunofluorescence staining of the V5 epitope (*51*). HEK293 cells were transfected with plasmids expressing DUX4-FL-V5 and CRISPR-Cas13b system components. The next day, cells were fixed in 4% paraformaldehyde (PFA) for 20-minutes, and nonspecific antigens blocked with 5% BSA in PBS supplemented with 0.2% Triton X-100. After overnight 4°C incubation in rabbit polyclonal anti-V5 primary antibody (1:2,500, Abcam, ab9116), cells were washed 3 times in PBS, and then incubated in goat anti-rabbit Alexa 594 secondary antibodies (1:2,500, Invitrogen) for 1 h at room temperature. After several PBS washes, cells were mounted using Vectashield mounting medium with DAPI (Vector Laboratories, Burlingame, CA).

### Quantitative Real-Time PCR (qRT-PCR) analysis of DUX4-activated biomarkers

To assess the specificity of the CRISPR/Cas13b system to target endogenous *DUX4* mRNA, 15Abic FSHD myoblasts (15A, 500,000 cells/reaction) were transfected with CRISPR/Cas13b and gRNA expression plasmids via electroporation as previously described (*51*). Subsequently, these cells were differentiated into myotubes for up to 7 days. Total RNA extraction was performed using TRIzol reagent (Thermo Fisher Scientific) following the manufacturer’s guidelines. Isolated RNA underwent DNase treatment (DNA-Free, Ambion, TX), followed by cDNA synthesis using a high-capacity cDNA reverse transcription kit (Applied Biosystems) with random hexamer primers. The resulting cDNA samples were used for TaqMan Assays with pre-designed primer/probe sets for *TRIM43*, *MBD3L2*, and *PRAMEF12* (markers of DUX4 activity), along with human *RPL13A* control primers/probes (Applied Biosystems). Normalization of all data was performed against non-targeting gRNA-transfected cells. This analysis was based on data from three to four independent experiments, each conducted in triplicate for every biomarker.

### RNA-Seq

We transfected 15A FSHD myoblasts with a 1:2 ratio of Cas13b and gRNA expression plasmids (mock transfection, Cas13b alone, or Cas13b with gRNAs 1, 2, 3, and 9). Three replicates were performed for each transfection. RNA was isolated using a MirVana kit (Thermo Fisher, AM1561) at 48 hrs, DNase treated, and sent to the Nationwide Children’s Hospital (NCH) Genomics Core for library construction and sequencing. Strand-specific RNA-seq libraries were prepared using NEBNext Ultra II Directional RNA Library Prep Kit for Illumina, following the manufacturer’s recommendations. In summary, total RNA quality was assessed using RNA 6000 Nano kit on Agilent 2100 Bioanalyzer (Agilent Biotechnologies) and concentration measured using Qubit RNA HS assay kit (Life Technologies). 500 ng aliquot of total RNA for each sample was rRNA depleted using NEB’s Human/Mouse/Rat RNAse-H based Depletion kit (New England BioLabs). Following rRNA removal, mRNA was fragmented and then used for first- and second-strand cDNA synthesis with random hexamer primers. ds-cDNA fragments underwent end-repair and a-tailing and were ligated to dual-unique adapters (Integrated DNA Technologies). Adaptor-ligated cDNA was amplified by limit-cycle PCR. Library quality was analyzed on Tapestation High-Sensitivity D1000 ScreenTape (Agilent Biotechonologies) and quantified by KAPA qPCR (KAPA BioSystems). Libraries were barcoded, pooled and sequenced at 2x150 bp read lengths on the Illumina HiSeq4000 platform to generate approximately 20 million paired-end reads per sample (range of 18M-24M). Adapters were trimmed with bcl2fastq2 (Illumina, v2.20).

#### Differential expression analysis

Reads were mapped to the GRCh38.13 human reference genome with GENCODE 32/Ensembl 98 gene annotations using STAR aligner (*120*) (2.7.0e, with options --twopassMode Basic and --outFilterMultimapNmax 50). A table of read counts per gene per sample was generated with featureCounts (*121*) (subread-2.0.0, with option -s 1 for strand-specific counting). Tests of differential gene expression were performed in R (4.3.2) using quasi-likelihood methods in the edgeR package (4.0.12) (*122*). Genes with low counts were filtered out, retaining the 19,834 genes with least 5 scaled counts in at least 3 samples, where the scaling compensates for differences in total counts between samples while maintaining the median of the total counts per sample. The function qlmQLFit (with non-default option prior.count=2) was used to fit the expression data to a model with a factor for “group” with six levels: Mock, Cas13b, gRNA1, gRNA2, gRNA3, and gRNA9, with three replicates in each group; group names denote, respectively, mock-transfected cells, cells transfected with Cas13b only (no guide RNA), and cells transfected with Cas13b in addition to the indicated guide RNA. Differential expression was assessed using the functions glmQLFTest and glmTreat with one contrast for each of the 15 possible pairwise comparisons between groups. glmQLFTest tests the hypothesis that the log2-fold-change (LFC) between groups in nonzero, and glmTreat tests the hypothesis that the LFC between groups has absolute value exceeding some cutoff, here taken to be 1 (corresponding to a two-fold change). In both cases, false-discovery rates (FDRs) (*123*) were computed from p-values to control for testing multiple genes. **Table 1** summarizes the number of differentially expressed (DE) genes for each contrast at three levels of stringency: (1) FDR < 0.05 for glmQLFTest; (2) FDR < 0.05 and abs(LFC)>1 for glmQLFTest; (3) FDR < 0.05 for glmTreat, which is more stringent than (2) since it required that abs(LFC) be *confidently* greater than 1 (as opposed to confidently greater than zero with a point estimate of at least 1). We use stringency level (2) in the discussion. Supplemental Table 2 includes results for all genes for each contrast.

#### Gene-set analysis

The R package goseq (1.54.0) (*124*) was used to perform gene set analysis while controlling for biases associated with gene lengths and expression levels by including logCPM as a covariate. The input DE genes were taken to be those with FDR < 0.05 for glmQLFTest and abs(LFC)>1, analyzed separately for up- and down-regulated genes, capped at a maximum of 500 genes in each direction (taking those with smallest p-value). The reference gene sets were taken from human Molecular Signatures Database (MSigDB, v2023.2.Hs) (*125*), restricted to sets containing between 3 and 500 genes. For conciseness, the discussion is limited to the collection of “hallmark” gene sets (*126*), with hits taken to be gene sets having nominal p-value for over-representation less than 10^-4^; this is not corrected for testing multiple gene-sets, but since the HALLMARK collection includes only 50 gene sets this cutoff gives family-wise error rate at most 0.005 by Bonferonni correction. The other collections of gene sets from MSigDB were tested as well, with results included in Supplemental Table 3.

#### Sequence-specific off-target analysis

The complementarity between each gRNA and each transcript was computed by aligning reverse-complemented gRNAs (rc-gRNAs) to transcripts. The reference transcriptome was constructed from the genome fasta and gene annotations (GTF) using the command rsem-prepare-reference from RSEM (1.3.0) (*127*). The function pairwiseAlignment in the R package Biostrings (2.70.3) (*128*) was run with type= “local” to find a best-scoring alignment of any subsequence of the rc-gRNA to any subsequence of each transcript, allowing mismatches and gaps (with match score = 1, mismatch score = -3, gap opening penalty = 4, gap extension penalty = 2). The score for each gene was defined to be the maximum score for any of the transcripts of the gene (based on Ensembl annotations). Scores for all genes and all gRNA are included in Supplemental Table 2. Because match score = 1, the local alignment score is at least as large as the longest contiguous stretch of perfect agreement. A cutoff of 11 on the local alignment score was used in the discussion section, as Cas13-induced knockdown of mRNA has been reported to be notably weaker when regions of complementary are 10 nucleotides or shorter (*69*).

### Animals

6- to 9-week-old C57BL/6 mice of both sexes were used in this study. All mice were housed in the vivarium of Nationwide Children’s Hospital (NCH), under specific pathogen-free conditions with a reversed 12-h light–dark cycle maintained at 23 °C with 30–70% humidity, and provided with food and automatic reverse osmosis watering. All cages are regularly changed and washed using 180-degree water and STERIS Cage-Klenz 180 and 200 detergents. This study complied with all relevant ethical regulations for animal testing and research. Animal studies followed the Care and Use of Laboratory Animals of the National Institutes of Health, and received ethical approval and were supervised by the NCH Institutional Animal Care and Use Committee (IACUC), approval number: AR13-00015.

The lead U6.gRNAs and Cas13b expression constructs were inserted into proviral AAV plasmids and used to manufacture AAV6 vectors using established protocols at Andelyn Biosciences, Columbus, Ohio. Mice were administered a total of 40 μl premixed viral vector cocktails containing 3x10^9^ DNAse resistant vector particles (DRP), also known as vector genomes (vg), of AAV6.CMV.DUX4-FL-V5; 7x10^10^ vg of AAV6.U6.gRNA; and 5x10^10^ vg of AAV6.miniCMV.Cas13b. As a control, contralateral co-injections were performed, either using 40 μl of saline or a premixed virus cocktail containing 3x10^9^ vg of AAV6.CMV.DUX4-FL-V5; 7x10^10^ AAV6.U6.nontar-gRNA, and 5x10^10^ vg of AAV6.miniCMV.Cas13b into the other TA. Muscles were harvested 3 weeks post-injection.

For analyzing Cas13b and gRNA expression at various time points and assessing the immune response against bacterial Cas13b, both TAs were injected only with 5x10^10^ vg of AAV6.miniCMV.Cas13b, 5x10^10^ vg of AAV6.U6.gRNA, 5 x 10^10^ vg of AAV6.mi405 (non-protein coding AAV) or saline. Muscles were collected 2-, 4-, 8-, and 24-weeks after injection, for qRT-PCR analysis of Cas13 and gRNA expression. To investigate the host immune response against AAV or the Cas13b protein, histological staining, flow cytometry, and fluorescent staining of various CD+ T-cells were performed 5 weeks after injection.

### Histological Analysis

TA muscles were dissected at indicated time points and flash frozen in O.C.T. Compound (Tissue-Tek) on liquid nitrogen-cooled isopentane. Ten-micrometer thick tissue sections were cut using a cryostat (Leica) and stored at -20°C until processed. Hematoxylin and eosin (H&E) staining was performed following standard protocols (*129*). The skeletal muscle sections were imaged using the Olympus BX61 microscope and analyzed with Olympus cellSens standard 1.9 software. Central nuclei quantifications were counted to assess muscle damage and regeneration (±SEM) (N=5 and N=7 per group).

### Droplet Digital PCR (ddPCR)

Please see methods for total RNA extraction and cDNA synthesis above. For ddPCR, a reaction mixture (20 µL) comprised of 1X ddPCR EvaGreen Supermix (Bio-Rad), 1 µM of forward and reverse DUX4 primers (identical to those utilized for qRT-PCR), and 1 µL of 10-fold diluted cDNA. We used the QX200 AutoDG Automatic Droplet Generator (Bio-Rad) to generate droplets, followed by amplification of the reactions in a C1000 Touch™ Thermal Cycler with a 96-Deep Well Reaction Module (Bio-Rad). The cycling parameters were defined according to the QX200 ddPCR EvaGreen Supermix guidelines. The annealing temperature was set at 60 °C based on the melting temperature (Tm) of the forward and reverse primers. Droplets were then read using the QX200 droplet reader (Bio-Rad). Data analysis was performed using the QuantaSoft analysis software 1.0 (Bio-Rad). For each 20 µL reaction, a minimum of 10,000 acceptable droplets was targeted for quantification. To quantify hsa-DUX4, mmu-Rpl13a, and mmu-Wfdc3 using TaqMan probes, the ddPCR reaction mixture (20 µL) included 1× ddPCR Supermix for probes (No dUTP), 1× specific TaqMan probe (ThermoFisher), and 50 ng synthesized cDNA. The cycling conditions were set as per the QX200 “ddPCR Supermix for probes” protocol with an annealing temperature of 58 °C.

### RNAscope assay and quantification

We used RNAscope *in situ* hybridization assay to measure *DUX4* and *Cas13b* mRNA levels in muscles co-injected with AAV6.CMV-DUX4, AAV6.miniCMV-Cas13b, and AAV6.U6-gRNA1 or AAV6.U6-nontar-gRNA control. Flash frozen muscle sections were cut at 10um thickness and kept frozen until processed. Tissue sections were fixed with 4% PFA for 30 min at room temperature, and dehydrated with 50%, 70%, and 100% ethyl alcohol gradients for 5 min each at room temperature. After rehydrating tissues with 70% and 50% ethyl alcohol gradients, RNAscope staining was performed as described previously (*23*) using custom-designed RNAscope probes specific to *DUX4* (Hs-DUX4-No-XMm, NM_001306068.2, ACDBio) or *Cas13b* (Probe-B-Psp-Cas13b-Codon, ACDBio).

### SYBR green-based quantitative RT-PCR gene expression assay for detecting gRNA and Cas13b

Total RNA was extracted using TRIzol reagent (Thermo Fisher Scientific) following the manufacturer’s protocol. After DNase treatment, cDNA was generated using Oligo(dT)18 primer provided in the Maxima H Minus First Strand cDNA Synthesis kit following the manufacturer’s instructions. The qPCR step was done using the SsoAdvanced Universal SYBR Green Supermix (Bio-Rad) according to manufactured protocol. Specifically, we mixed 1× of SsoAdvanced SYBR Green Supermix with 50 ng of cDNA, and with 1 µM of forward and reverse primers (Supplementary Table 2). The DNA polymerase activation and DNA denaturation were carried out at 95 °C for 30 seconds. The PCR step included 39 cycles of denaturation step at 95 °C for 10 seconds, followed by an annealing/extension and plate reading step at 60 °C for 30 seconds. The amplification step was completed with a melt-curve analysis step (snap cool to 3 °C followed by an increment of 0.5 °C every 5 s until reaching 95 °C). The qRT-PCR analyses were performed using the Bio-Rad CFX96 Real-time system.

### Tissue Immunofluorescence Staining

10 um thick muscle sections fixed in 4% PFA and incubated with 1:500 rabbit anti-V5 (Abcam, ab9116), or 1:500 Rabbit anti-Cleaved Caspase-3 (Cell Signaling Tech, #9664), or anti-CD3 (Abcam, ab16669), CD4 (Abcam, ab183685), CD8 (Abcam, ab217344) following a previously published protocol (*14*). Fluorescence images of skeletal muscle sections were taken using the Olympus BX61 microscope and the Olympus cellSens standard 1.9 software.

### Flow Cytometry Analysis

*Live and dead cell discrimination.* Cells were suspended in 100 μl of Zombie NIR viability dye (1:1000 in 1X PBS) for 30 minutes on ice while protected from light. Fc receptor blocking of muscle single-cell suspensions was performed by incubation with an anti-CD16/32 antibody (clone 2.4G2) before staining. Single-cell suspensions were stained with a panel of antibodies against several cell surface antigens (**Table S4**). Analysis was performed on live cells on a BD LSRFortessa™ X-20 Cell Analyzer with FACSDiva software (BD Bioscience). Post-acquisition analysis was performed with Flowjo software version 10.8.

### Statistical analysis

Statistical analyses for caspase-3/7 assay, cell viability assay, RNAscope, western blot, and qRT-PCR data were performed in GraphPad Prism Software version 9 (GraphPad, La Jolla, CA) using the indicated statistical tests. Statistical significance was defined as P-value < 0.05 unless otherwise noted, and number of replicates *n* is indicated in the figure legends. Data are expressed as mean ± SEM unless otherwise noted. All animals and data points were included in the analyses without any exclusions.

Statistical analysis of RNA-seq data is described in the RNA-seq methods section.

## Acknowledgments

This work was supported by an investigator-initiated grant from Friends of FSH Research to A.R., FSHD Society postdoctoral fellowship to A.R., and National Institute of Arthritis and Musculoskeletal and Skin Diseases (NIAMS) 1R01AR062123-05 to S.Q.H. During the course of this study, G.A.C.’s salary was supported by a post-doctoral grant from the FSHD Society (grant 5888746560). The authors thank Dr. Charles Emerson and the University of Massachusetts Wellstone Muscular Dystrophy Program for providing the human immortalized myoblasts used in our studies. We thank Amy Wetzel from the Biomedical Genomics Core of the Research Institute at Nationwide Children’s Hospital for help with RNA-seq data. The schematic dual-reporter screening of collateral activity included as part of Figure 3A is created in BioRender (Rashnonejad, A. (2025) https://BioRender.com/w66y309).

## Author contributions

A.R. conceptualized and obtained funding for the project, designed and cloned CRISPR-Cas13b gRNAs and other plasmids in this study, performed or directed all experiments, created figures and legends, and wrote/edited the paper. G.A.C., injected and dissected animals, contributed to central nuclei counts, performed the RNAscope assay and quantified the corresponding results. N.K.T. cloned dual-reporter and AAV.dGFP/DUX4 plasmids, contributed to western blots and central nuclei count. M.A. contributed to writing and organizing the manuscript, G.C. and A.V. performed flow cytometry and quantified the corresponding results. A.F. performed AAV vector production, O.D.K. performed RNAseq data analysis, made figures, and participated in manuscript writing and editing. S.Q.H. was responsible for study design contributions, obtaining funding for the project, establishing collaborations to complete the work, and writing/editing the manuscript.

---

**Table S1.**
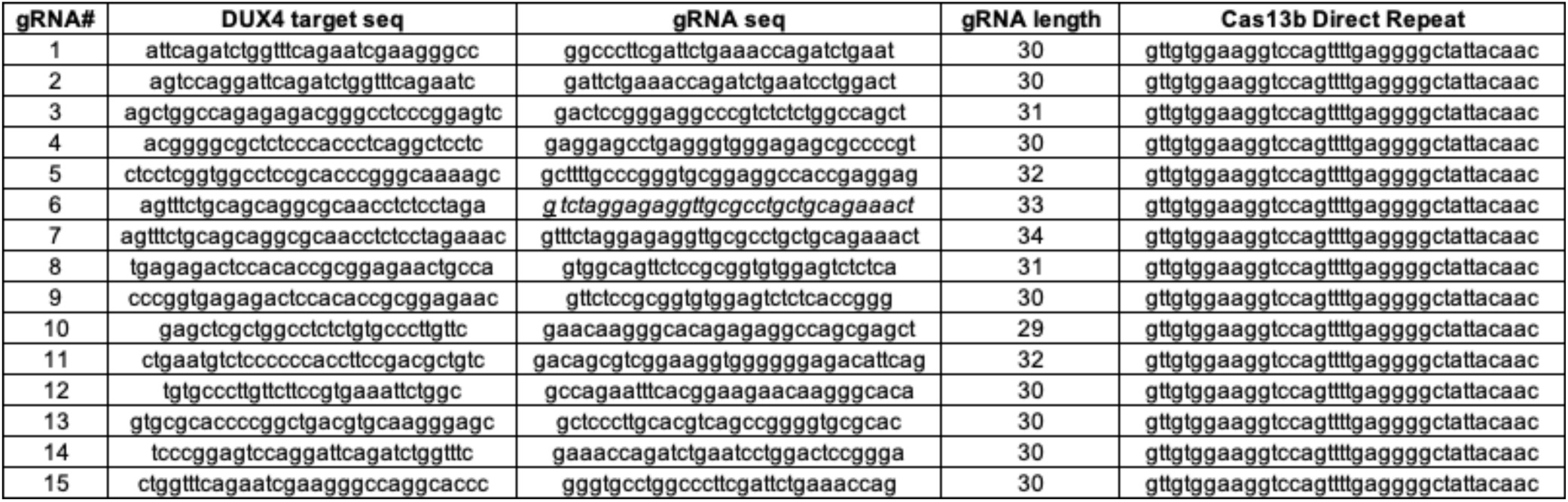
Cas13b gRNA sequences and target sites.

**Table S2.** RNA-seq differential gene expression results. This table includes differential expression analysis results for all genes (after filtering out genes with very low expression) for all 15 contrasts between pairs of conditions. The prefix of column names indicates the contrast, e.g. gRNA1_vs_Cas13b for the contrast of Cas13b with gRNA1 versus Cas13b alone. Column-names ending PValue and FDR are for glmQLFTest and column-names ending PValue2 and FDR2 are for glmTreat with lfc=1 (see RNA-seq Methods). The column logCPM gives log_2_(counts-per-million) averaged across all groups. The columns local_gRNAn for n = 1, 2, 3 or 9 give the maximum local alignment score between the reverse complement of gRNAn and any transcript of the gene (see RNA-seq methods). The table also includes columns with raw read counts for each gene in each sample.

(Please see additional supplementary materials)

**Table S3.** RNA-seq gene-set analysis results. This Excel spreadsheet includes the full results for the gene set analyses described in the RNA-seq methods section. There are separate sheets for each of the 15 between-group comparisons, and each sheet includes results for 27 MSigDB gene-set collections or subcollection, further divided into separate analyses of the DEGs that were up and down in the between-group comparisons. The first five columns, taken directly from the output of the function goseq from the R package of the same name, give the category name, the p-value for this category being over-represented among DEGs, the p-value for this category being under-represented among DEGs genes, the number of DEGs in the category, and the total number of genes in the category. These have been augmented with columns named collection (MSigBD collection or subcollection); direction (up or down); adjusted.p.value (using FDR, separately for combination of collection or subcollection and direction); nGenes (number of genes in universe after filtering and restricting to genes with at least one annotation in the collection or subcollection being analyzed); nDE (number of input DEGs after filtering); and genesDEInCat (comma-separated list of gene symbols for DEGs in category, up to a maximum of 100 gene symbols per category, and included only for the top 100 categories ranked by over_represented_pvalue and with a cutoff of over_represented_pvalue < 0.01).

(Please see additional supplementary materials)

**Fig S1.**
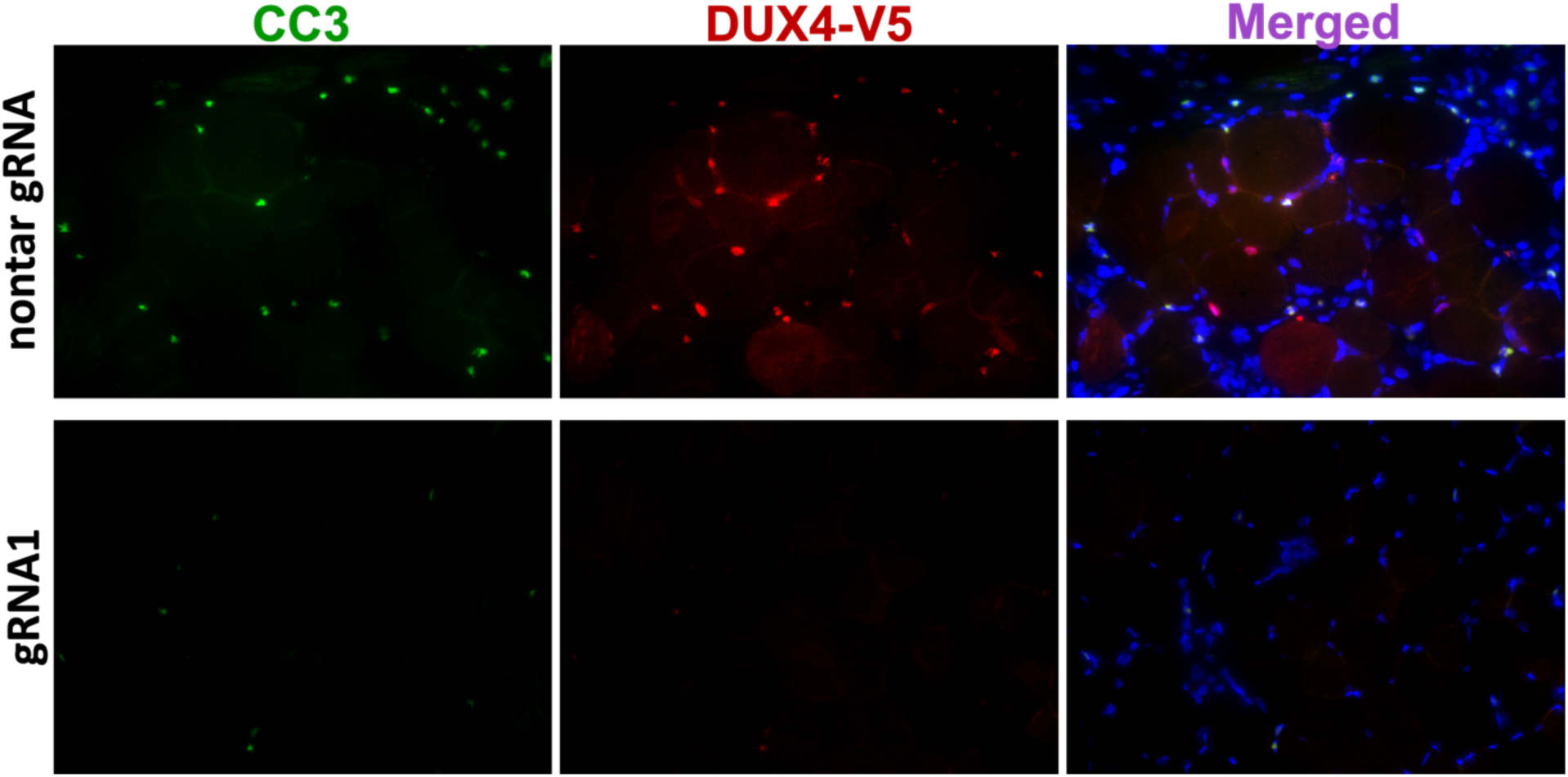
Apoptotic nuclei were reduced in treated muscles. Immunofluorescent staining showed reduced DUX4-V5 (red) and Cleaved Caspase3 (CC3, green) signals in CRISPR-Cas13b treated FSHD muscles.

**Table S4.** The list of antibodies used for flow cytometry analysis.

| Marker | Color | Vendor | Cat # | Clone | Final dilution |
| --- | --- | --- | --- | --- | --- |
| Viability | Zombie NIR | BioLegend |  |  | 1:1000 |
| CD3 | ef-450 | ebio | 48-0032-82 | 17A2 | 1:100 |
| CD4 | BV605 | BioLegend | 100548 | RM4-5 | 1:500 |
| CD8 | FITC | BioLegend | 100706 | 53-6.7 | 1:500 |
| CD44 | PE | BioLegend | 103024 | IM7 | 1:1000 |
| CD62L | APC | BioLegend | 104412 | MEL-14 | 1:100 |
| CD69 | PECy7 | BioLegend | 104512 | H1.2F3 | 1:500 |
| CD107a | PerCP-Cy5.5 | BioLegend | 121625 | 1D4B | 1:100 |
| CD45 | BUV395 | BD biosciences | 564279 | 30-F11 | 1:500 |
| CD90.2 | BV785 | BioLegend | 105331 | 30-H12 | 1:100 |

